# Metabolomic analysis of Chinese chestnut (*Castanea mollissima* Bl.) leaves in response to *Oligonychus ununguis* feeding stress

**DOI:** 10.1101/2025.11.20.689536

**Authors:** Xinfang Zhang, Shuhang Zhang, Ying Li, Yan Guo, Jinyu Liu, Jiayi Liu, Liying Fan, Guangpeng Wang

**Affiliations:** Changli Research Institute of Fruit Trees, Hebei Academy of Agricultural and Forestry Sciences, Changli, Hebei, 066600, China

**Keywords:** *Castanea mollissima* Bl., *Oligonychus ununguis*, secondary metabolites, metabolomics

## Abstract

Piercing-sucking insects such as the chestnut red mite (CRM; *Oligonychus ununguis*) damage leaves by extracting mesophyll cell nutrients, thereby reducing photosynthesis and nut yield in Chinese chestnut (*Castanea mollissima* Blume). However, the metabolic basis of the chestnut response to mite stress remains elusive. To address this, leaf metabolomics was conducted on four varieties with different resistance levels. Resistance grades were determined by the leaf chlorosis index, and three comparison groups were established. Secondary metabolome profiling revealed that resistant varieties showed greater metabolic changes under infestation than susceptible ones. In total, 713 total metabolites and 275 metabolites with significantly differential accumulation among germplasms were identified across the comparisons. KEGG enrichment highlighted 246 metabolites and 75 differential metabolites involved in flavonoid, flavonol, and phenylpropanoid biosynthesis. Notably, 27 phenylpropanoids and flavonoids accumulated significantly in resistant varieties but were largely absent or unchanged in susceptible ones. These findings suggest that specialized phenylpropanoids and flavonoids, along with their associated pathways, play key roles in CRM resistance, providing a foundation for breeding mite-resistant chestnut varieties.

## Introduction

Spider mites are globally significant pests that reduce yields in crops and fruit trees, leading to major economic losses^[1]^. Their rapid evolution, driven by heavy pesticide use, has resulted in resistance to several chemical groups^[2-3]^. As a result, improving mite resistance in fruit tree breeding is a central objective for sustainable agriculture^[4]^.

The Chinese chestnut (*Castanea mollissima* Blume), an ancient domesticated species of the Fagaceae family, originates from China, which remains the world’s leading chestnut producer. This species is valued for its nut quality and strong tolerance to environmental stresses^[5]^. In northern regions such as Shandong, Hebei, and Beijing, chestnut red mite (*Oligonychus ununguis*) is a major pest. Its feeding causes chlorosis along leaf veins, severely impairing photosynthesis and lowering both nut yield and quality during the infestation year and the following season^[6]^. For growers, this results in serious economic losses, with mite control accounting for 20-25% of total production costs. Breeding and cultivating mite-resistant chestnut varieties therefore represent the most sustainable way to reduce inputs and promote environmentally sound production.

Although current statistics are incomplete, plants are estimated to produce between 200,000 and one million unique metabolites^[7-8]^. Differences in metabolite diversity and abundance reflect plant adaptability to environmental stress. Secondary metabolite synthesis is a central strategy for resisting both biotic and abiotic challenges^[9]^. In response to insect or mite attack, plants employ constitutive and induced defenses, including growth inhibition and metabolic adjustments^[10-11]^. Considerable progress has been made in elucidating these metabolic responses. For example, in peach, resistance to the *Myzus persicae* is associated with the production of triterpenoids and upregulation of their biosynthesis-related genes^[12]^. In maize, feeding by Asian corn borer (*Ostrinia furnacalis*) enhances an array of defenses linked to phytohormones, benzoxazinoids, and volatiles during the mid-whorl stage under field conditions^[13]^. In tea, infestations by *Oligonychus cofeae* Nietner and *Helopeltis theivora* Waterhouse stimulate biosynthesis of fatty acid derivatives^[14]^, while catechins contribute to horse chestnut resistance against leaf miner^[15]^. Combined transcriptomic and metabolic analyses further suggest that MAPK and Ca^2+^-mediated PR-1 signaling, as well as flavonoid and phenylpropanoid biosynthesis, play roles in tea plant responses to the six-spotted spider mite^[16]^.

Few studies have systematically examined the accumulation and dynamic changes of secondary metabolites in Chinese chestnut under mite stress. The differences in metabolite profiles and associated pathways between resistant and susceptible varieties also remain poorly understood. In recent years, widely targeted metabolomics, primarily using ultra-high performance liquid chromatography alongside triple quadrupole mass spectrometry (UHPLC-QQQ-MS), has emerged as a robust technique that integrates the strengths of non-targeted metabolic analysis^[17]^. Owing to its high throughput, broad coverage, rapid separation, and sensitivity, this approach has been successfully utilized for metabolite studies in crops and fruit trees^[18]^. It therefore provides an effective strategy for qualitative and quantitative analyses of secondary metabolism in Chinese chestnut tissues.

Although metabolic defense mechanisms against piercing-sucking pests have been widely characterized in model plants, comparable research in woody species, particularly economic trees such as Chinese chestnut, remains limited. In this study, four *C. mollissima* varieties with differing resistance to *O. ununguis* were analyzed: Yanxing (S19), Xujia 1 (R-1), Yankui (M1), and Yanshanzaofeng (M2). Secondary metabolite profiles and relative contents in mite-infested leaves were examined using a broadly targeted metabolomics program. Comparative analysis identified differential metabolites and pathways between resistant and susceptible varieties. These findings advance understanding of chestnut leaf responses to *O. ununguis* stress and provide a basis for breeding new mite-resistant cultivars.

## 3. Results

### 3.0 Classification of mite resistance in Chinese chestnut germplasms

Mite resistance grades in the four germplasms (S19, R-1, M1, and M2) was classified using the leaf chlorosis index. S19 showed a chlorosis index of 68.73%, and was classified as highly susceptible, while R-1 had an index of 12.35% and was classified as highly resistant. M1 (45.38%) and M2 (37.24%) were both categorized as susceptible. Based on these resistance levels, three comparison groups were established: R-1 vs. S19, R-1 vs. M1, and R-1 vs. M2. These pairings represent either extreme differences (high resistance vs. high susceptibility) or moderate differences (high resistance vs. susceptibility), enabling systematic analysis of secondary metabolite variation and changes in key pathways across resistance gradients.

### 3.1 Metabolite identification and quality control assessment

Total ion current (TIC) plots and multi-peak detection plots of a quality control (QC) sample are shown in Figure 1. The TIC plot provides a continuous profile of summed ion intensities over time, while the multiple reaction monitoring (MRM) multi-peak plot illustrates the presence of multiple metabolites, with each colored peak representing a distinct compound. Using a local metabolite database, both qualitative and quantitative MS were performed. In total, 713 metabolites were identified: 83 flavones, 100 flavonols, 38 flavonoids, 15 flavanols, 10 isoflavones, 221 phenolic acids, 43 tannins, 58 alkaloids, 72 terpenoids, 31 lignans, 18 coumarins, one steroid, and 23 other metabolites. Full details are provided in Table S1.

**Figure 1.**
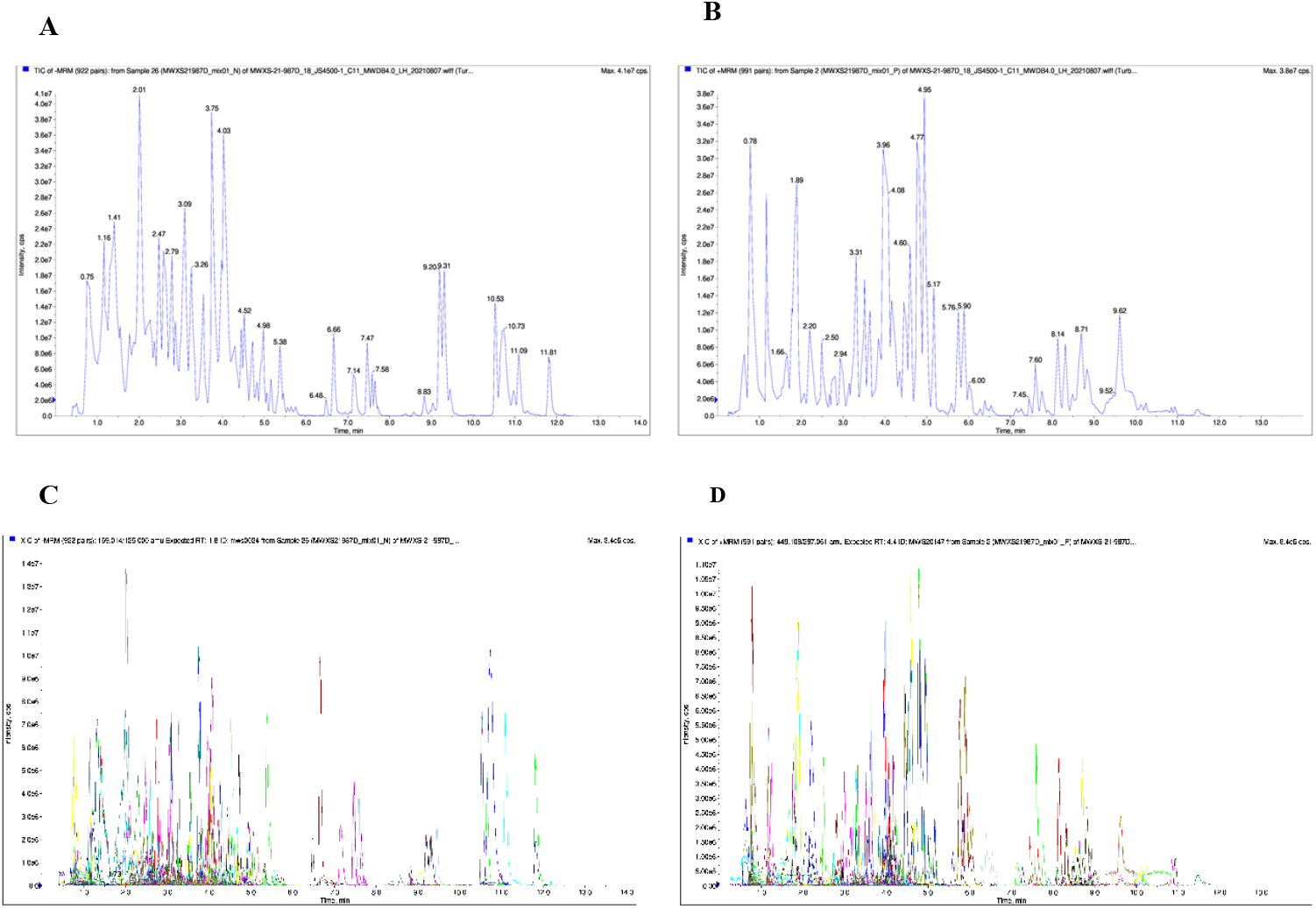
(**A**) Total ion current diagrams of QC samples: negative ion mode. (**B**) Total ion current diagrams of QC samples: positive ion mode. (**C**) Multi-peak MRM plots: negative ion mode. (**D**) Multi-peak MRM plots: positive ion mode.

Overlay analysis was used to assess the technical repeatability of compound extraction and detection. As shown in Figure 1, TIC plots from three QC samples overlapped almost perfectly, indicating stable and repeatable detection across time points. In addition, strong correlation coefficients (r = 0.989–0.998) among each sample’s three biological replicates confirmed acceptable homogeneity (Fig. S2).

### 3.2 Multivariate analysis of metabolite profiles in Chestnut chestnut germplasms

A heatmap was generated to assess correlations among the three biological replicates, with Pearson correlation coefficient (PCC) as the evaluation criterion. The results showed strong positive correlations across all replicates (Fig. 2A).

**Figure 2.**
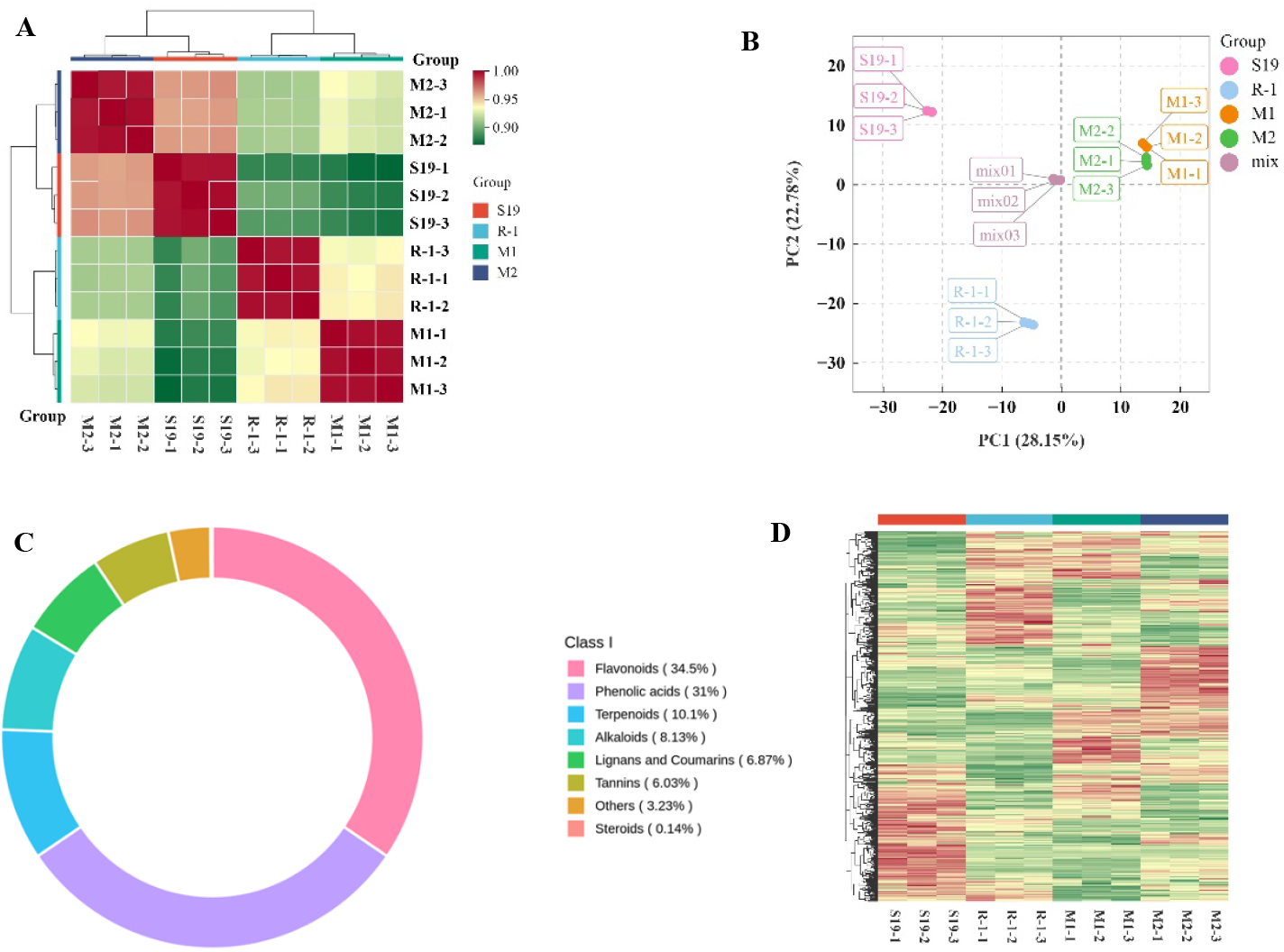
(**A**) Heatmap showing correlations among the three biological replicates of the four Chinese chestnut varieties. (**B**) PCA score plot of metabolite profiles. (**C**) Circular diagram of metabolite categories detected. (**D**) Clustering heat map of four varieties.

Principal component analysis (PCA) was performed using LC-MS/MS with electrospray ionization (LC-ESI-MS/MS) to compare overall metabolic differences among the four chestnut germplasms^[19]^. In the PCA score plot, PC1 and PC2 accounted for 28.15% and 22.78% of the variance, respectively. The results showed minimal differences between M1 and M2, while R-1 separated clearly from the other samples along the PC2 axis, indicating distinct metabolite profiles between resistant and susceptible germplasms. The tight clustering of three biological replicates for each variety further demonstrated experimental reproducibility and reliability (Fig. 2B).

In total, 713 secondary metabolites were identified in leaves of the four chestnut varieties: 246 flavonoids, 221 phenolic acids, 72 terpenoids, 58 alkaloids, 49 lignins and coumarins, 43 tannins, 23 other compounds, and one steroid. Flavonoids and phenolic acids were the dominant classes, comprising 34.5% and 31% of the total, respectively, while steroids and other compounds were minor components, accounting for only 0.14% and 3.23% (Fig. 2C).

Hierarchical cluster analysis (HCA) of the bioactive compounds separated the samples into four distinct classes on the heatmap. This indicated significant differences in metabolite abundance among the chestnut varieties, suggesting that genetic variation strongly shapes metabolite profiles associated with red mite resistance (Fig. 2D).

### 3.3 Discriminant analysis of metabolite profiles in response to mite stress

As a supervised method, orthogonal partial least squares-discriminant analysis (OPLS-DA) combines PLS-DA with orthogonal signal correction (OSC), decomposing the X matrix into variation related to Y and unrelated variation^[20]^. Differentially responsive variables are then identified by excluding the irrelevant components. In this study, OPLS-DA was utilized to identify metabolic differences between pairs of resistant and susceptible varieties: M1 vs. R-1 (R^2^Y = 1, R^2^X = 0.751, Q^2^ = 0.986), R-1 vs. M2 (R^2^Y = 1, R^2^X = 0.743, Q^2^ = 0.984), and S19 vs. R-1 (R^2^Y = 1, R^2^X = 0.751, Q^2^ = 0.986). All models showed R^2^X values above 0.55, R^2^Y values above 0.99, and Q^2^ values above 0.9, confirming their stability. The OPLS-DA score plots revealed clear pairwise separation, indicating distinct metabolite responses to mite stress between resistant and susceptible varieties. Corresponding S-plots are shown in Figure 3D-F. model reliability was further validated using 200 random alignment and permutation tests.

**Figure 3.**
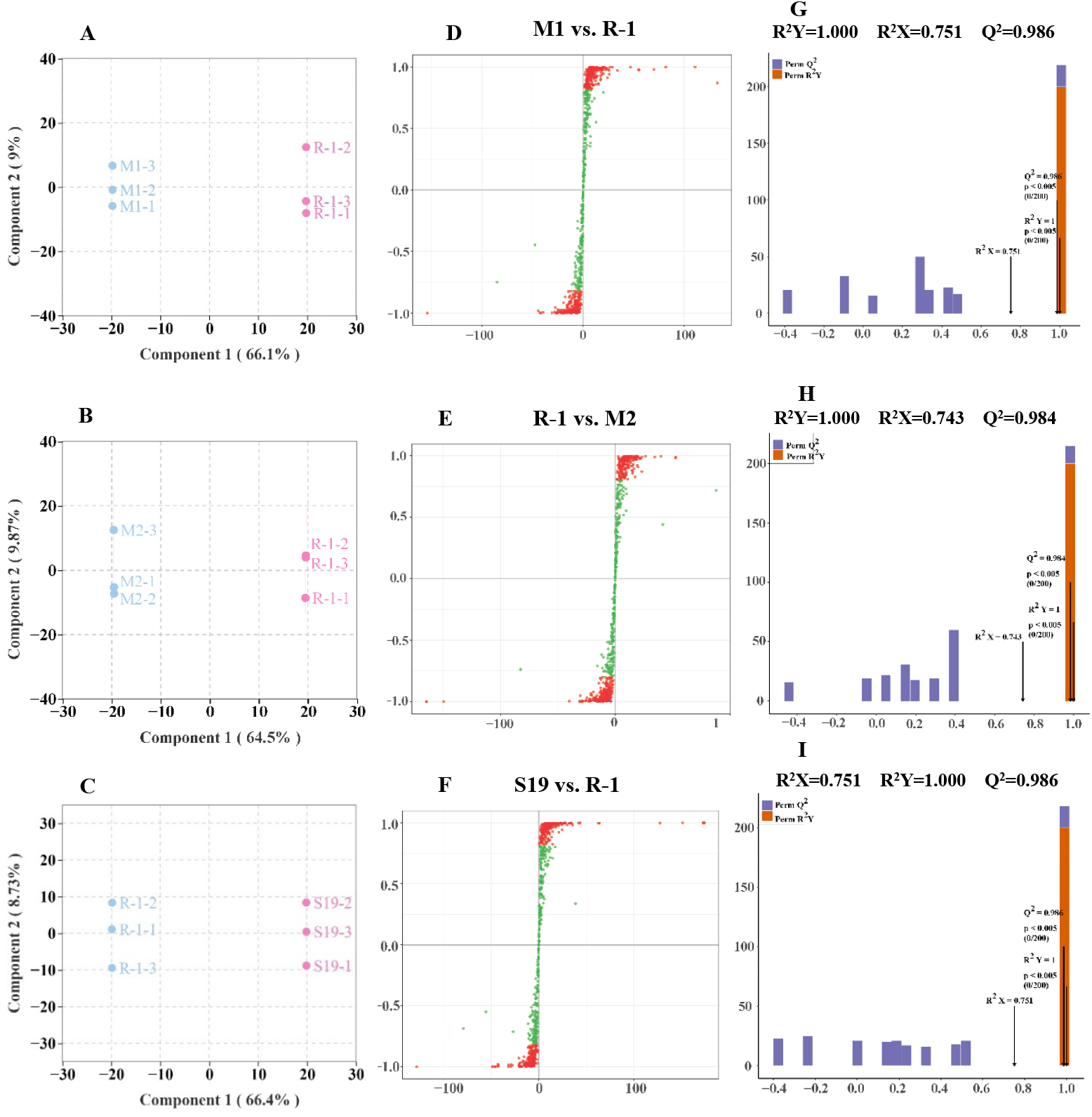
OPLS-DA score plots of four Chinese chestnut germplasms. (**A**) M1 vs. R-1, (**B**) R-1 vs. M2, and (**C**) S19 vs. R-1.

### 3.4 Differential metabolite discovery and pathway enrichment analysis

Differential metabolites were discovered across all three pairwise comparisons by utilizing the variable importance in projection (VIP) ≥ 1 as well as fold change (FC) ≤ 0.5 or ≥ 2 or thresholds. In total, 128 significantly differential metabolites were found in M1 vs. R-1 (63 upregulated, 65 downregulated), 133 in R-1 vs. M2 (56 upregulated, 77 downregulated), and 118 in S19 vs. R-1 (45 upregulated, 73 downregulated). Volcano plots of these comparisons are shown in Figure 4A–F, and the top-ranked metabolites by log_2_FC are presented in Figure S3.

**Figure 4.**
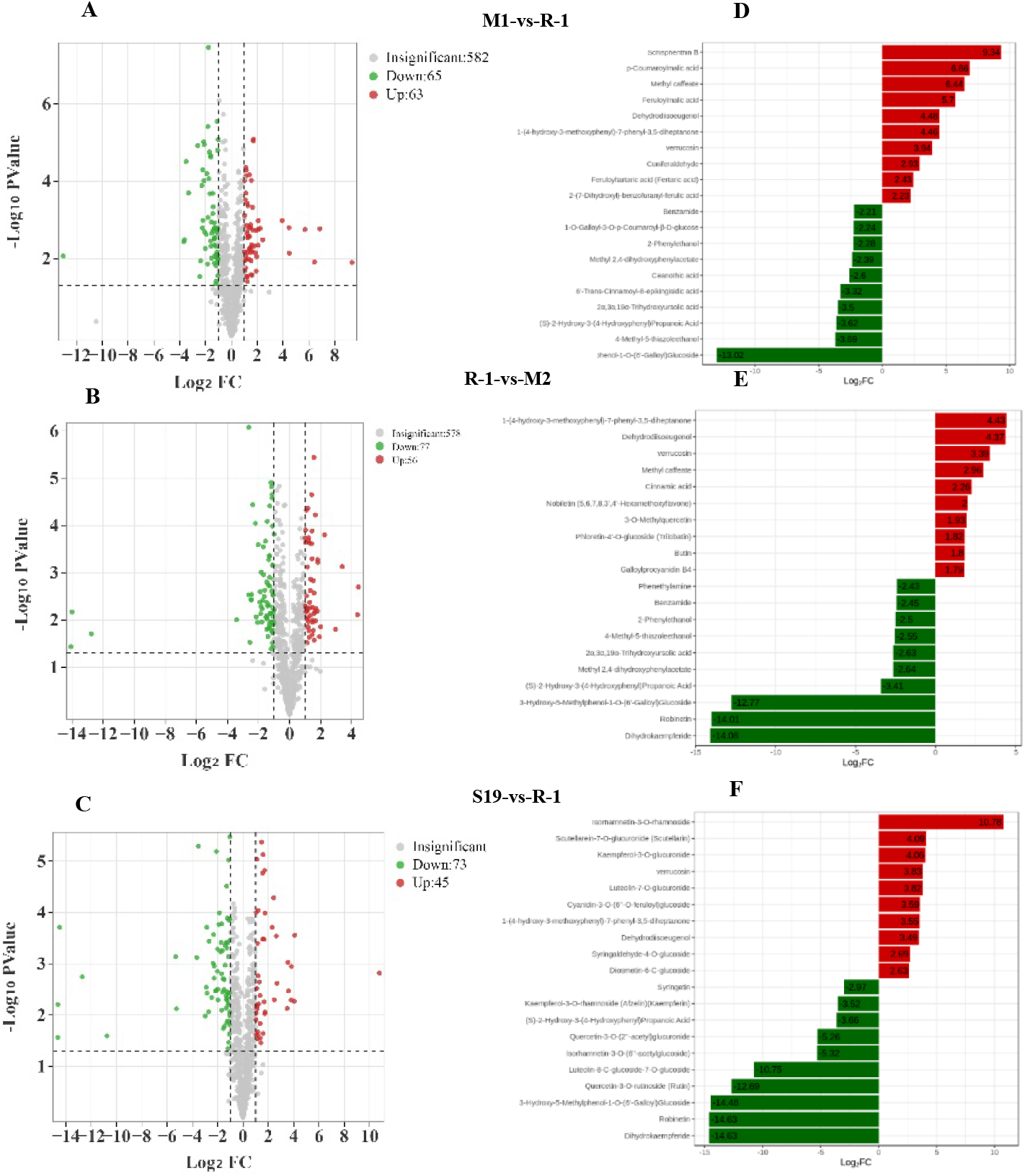
Volcano plots of significantly differential metabolites in different pairwise comparisons: (**A**) M1 vs. R-1, (**B**) R-1 vs. M2, and (**C**) S19 vs. R-1. Bar charts of fold-change (FC) differences: (**D**) M1 vs. R-1, (**E**) R-1 vs. M2, and (**F**) S19 vs. R-1. Red indicates upregulation and green indicates downregulation.

Significantly differential metabolites from each pairwise comparison were mapped to the Kyoto Encyclopedia of Genes and Genomes (KEGG) database, a centralized resource for metabolic pathway analysis linking gene expression and metabolite accumulation. Pathway enrichment analysis classified the differential metabolites into distinct pathways. Comparisons of the resistant variety R-1 with the susceptible varieties M1, M2, and S19 involved 14, 20, and 13 pathways, with the major pathways illustrated in bubble plots (Figure 5).

**Figure 5.**
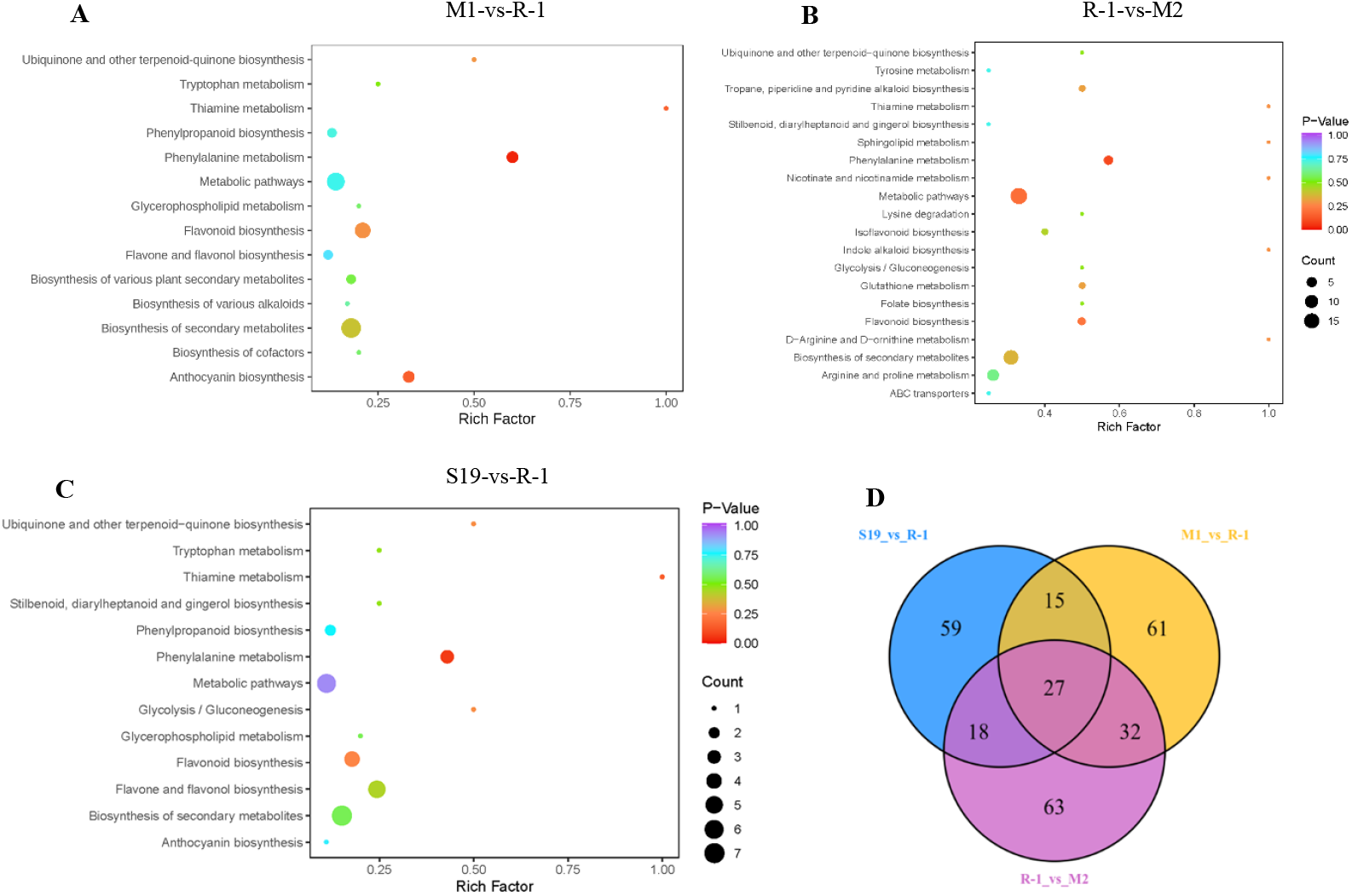
KEGG enrichment maps of differential compounds identified among pairwise comparisons: (**A**) M1 vs. R-1, (**B**) R-1 vs. M2, and (**C**) S19 vs. R-1. (**D**) Venn diagram of differential metabolites across the three comparisons.

Between M1 and R-1, differential compounds were significantly enriched in phenylalanine metabolism as well as flavonoid and anthocyanin biosynthesis. Flavonoid biosynthesis and secondary metabolite biosynthesis contained the highest numbers of differential metabolites, highlighting their central role in metabolic differences between the groups. In R-1 vs. M2, enrichment was observed in phenylalanine metabolism, general metabolic pathways, and flavonoid biosynthesis, with additional metabolites mapped to secondary metabolite biosynthesis as well as arginine and proline metabolism. For S19 vs. R-1, significant enrichment also occurred in phenylalanine metabolism and flavonoid biosynthesis, with many metabolites assigned to general metabolic pathways as well as flavonoid and secondary metabolite biosynthesis.

A total of 275 differential metabolites were identified across the comparisons of R-1 with S19, M1, and M2, including 27 common metabolites revealed by Venn analysis (Figure 4, Table 1). These were categorized into eight bioactive groups: steroids, terpenoids, alkaloids, tannins, lignans and coumarins, flavonoids, phenolic acids, and other compounds (Table 2). The 27 common metabolites showed consistent expression patterns across all groups and included phenolic acids, flavonoids, alkaloids, and terpenoids. Six compounds (cinnamic acid, methyl caffeate, baicalein, gallocatechin, dehydrodiisoeugenol, and lucidenic acid) accumulated specifically at high levels in the resistant variety. Among the 27 common metabolites, 15 showed highly significant differences (*P* < 0.01), with 11 downregulated and four upregulated. KEGG annotation identified five metabolites (2-phenylethylamine, 2-phenylethanol, 4-methyl-5-thiazolylethanol, cinnamic acid, and gallocatechin), which were mainly enriched in stress-related pathways including flavonoid metabolism (map00941), phenylalanine metabolism (map00360), and phenylpropanoid biosynthesis (map00940).

**Table 1.**
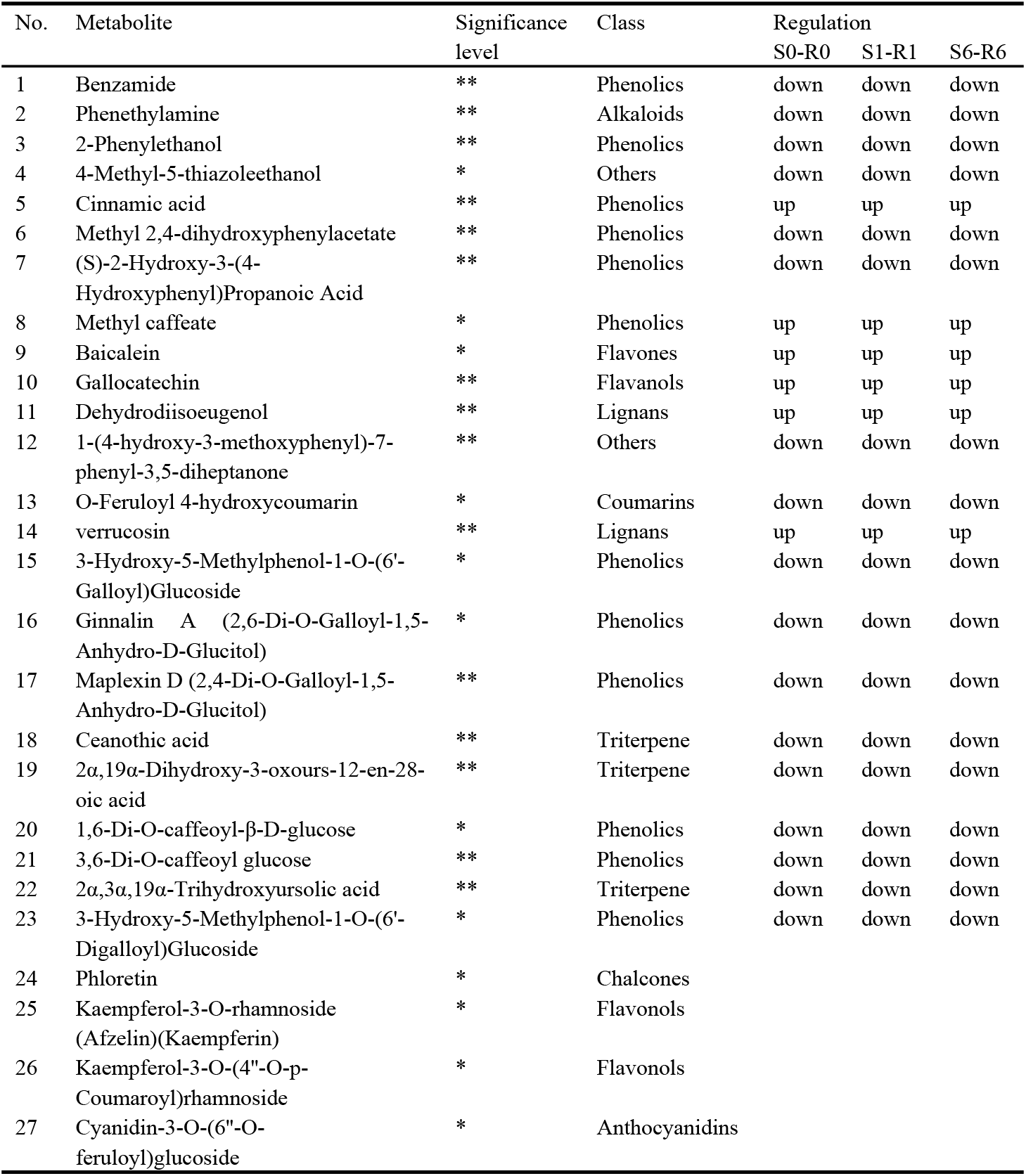
Twenty-seven differential metabolites common to all comparison groups.

**Table 2.** Number of differential compounds in *Castanea mollissima* leaves with varying levels of mite resistance.

| Group Class | S19 vs. R-1 |  | M1 vs. R-1 |  | M2 vs. R-1 |  |
| --- | --- | --- | --- | --- | --- | --- |
|  | Up | Down | Up | Down | Up | Down |
| Others | 1 | 1 | 3 | 2 | 3 | 3 |
| Steroids | 0 | 1 | 0 | 0 | 0 | 1 |
| Terpenoids | 1 | 4 | 1 | 19 | 4 | 17 |
| Alkaloids | 4 | 1 | 7 | 3 | 1 | 10 |
| Tannins | 0 | 4 | 7 | 1 | 8 | 1 |
| Lignans and Coumarins | 10 | 4 | 6 | 3 | 5 | 6 |
| Flavonoids | 19 | 29 | 23 | 17 | 29 | 14 |
| Phenolic acids | 10 | 30 | 18 | 25 | 11 | 27 |

## 2. Materials and methods

### 2.1 Plant materials

Four Chinese chestnut varieties (Yanxing, Xujia 1, Yankui, and Yanshanzaofeng) were planted in 2021 at the Chestnut Germplasm Resource Nursery of the Changli Institute of Pomology, Hebei Academy of Agricultural and Forestry Sciences, Qinhuangdao, Hebei Province, China (39°72′ N, 119°15′E). Leaves from the middle of vegetative branches were collected. Insects were removed prior to storing the samples at -80°C in an ultra-low temperature unit.

### 2.2 Field evaluation of mite resistance in Chinese chestnut varieties

Field investigations were conducted to evaluate mite resistance following the descriptors and data standards for *C. mollissima*^[21]^. For each variety, ten trees of similar growth were selected: five survey trees and five controls. Survey trees received no pest control during the growing season, while control trees were sprayed with pesticides at the occurrence period of chestnut red spider mite to strictly limit damage. In early August, chlorotic areas were measured on 50 leaves per tree, using the grading standard shown in Table 3.

**Table 3.** Grading standard for chestnut leaf chlorosis after mite damage.

| Standard score | Description |
| --- | --- |
| 0 | Leaf blade is almost entirely green |
| 1 | Chlorotic area < 25% of the total leaf area |
| 2 | Chlorotic area 26-50% of the total leaf area |
| 3 | Chlorotic area 51-75% of the total leaf area |
| 4 | Chlorotic area 76-100% of the total leaf area |

The leaf chlorosis index was calculated based on the grading standard using the formula:

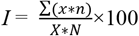

where *I* signifies the leaf chlorosis index, *x* signifies the chlorosis grade value, *n* signifies the number of leaves at that grade, *X* signifies the highest grade value, and *N* signifies the total number of leaves. Resistance to red mite was classified as: highly resistant (*I* < 15%), resistant (15% ≤ *I* <25%), moderately resistant (25% ≤ *I* <35%), sensitive (35% ≤ *I* <60%), or highly sensitive (*I* ≥ 60%).

### 2.3 Leaf sample collection for metabolomic analysis

Based on phenotypic data of mite resistance in *C. mollissima*, leaf samples were collected concurrently with the field investigation. Each biological replicate included nine plants exposed to mite damage. From peripheral vegetative branches, 20 leaves per replicate were pooled, inspected to remove impurities (insects or debris), flash-frozen in liquid nitrogen, and reserved at −80°C.

### 2.4 Sample preparation for metabolomic analysis

Freeze-dried samples were ground in a mixer mill (MM 400, Retsch, Haan, Germany) at 30 Hz for 1.5 min. Approximately 100 mg of each powdered sample was extracted with 1.2 mL of 70% methanol (aqueous) overnight at 4°C, followed by centrifugation at 12,000 g for 10 min. The supernatant was purified using a CNWBOND Carbon-GCB SPE Cartridge (250 mg, 3 mL; ANPEL, Shanghai, China), filtered through a 0.22 μm membrane (SCAA-104; ANPEL, Shanghai, China), and reserved in glass vials for downstream analyses. QC samples were made from pooled four varieties extracts using the same procedure.

### 2.5 Analytical setup for metabolomic profiling

LC-MS analysis was performed using a LC-ESI-MS/MS system (HPLC: Shim-pack UFLC, Shimadzu CBM30A; MS: Applied Biosystems 4500 Q TRAP, Framingham, MA, USA). Separation was achieved on a Waters ACQUITY UPLC HSS T3 C18 column (1.8 µm, 2.1 mm × 100 mm). The mobile phases included 0.04% acetic acid in water (solvent A) and 0.04% acetic acid in acetonitrile (solvent B). The gradient began at 95% A and 5% B, shifted linearly to 5% A and 95% B within 10 min, held for 1 min, and then returned back to 95% A and 5% B within 0.1 min and held for 2.9 min. The column oven was kept at 40°C, and the injection volume was 4 μL. The ion spray voltage was set to 5500 V (positive ion mode) and -4500 V (negative ion mode), with a source temperature of 550°C. Ion source gasses I (GSI) and II (GSII) were respectively set to 50 and 60 psi; curtain gas (CUR) to 30 psi; and collision gas (CAD) to high. In QQQ mode, nitrogen was used as the CAD at 5 psi.

### 2.6 Qualitative and quantitative analysis of metabolites

Metabolites were identified and quantified using MS with both a public database and self-built MetWare Database (Wuhan MetWare Biotechnology Co., Ltd., Wuhan, China). Quantification was performed in MRM mode on a QQQ MS^[22]^. In this mode, characteristic ions of each metabolite were filtered to generate signal intensities, and chromatographic peaks were integrated using MultiQuant software. The peak area of each chromatographic peak reflected the relative content of the corresponding metabolite.

### 2.7 Evaluation of analytical accuracy and reproducibility

To assess repeatability under mite stress, QC samples (pooled extracts) were analyzed following each group of 10 experimental samples^[23]^. Accuracy and reproducibility were evaluated by overlay analysis of MS TIC profiles from the QC samples.

### 2.7 Multivariate and statistical analyses

Raw data were normalized as described previously^[24]^ and log_2_ transformed before analysis. PCA, HCA, and OPLS-DA were carried out in R (http://www.r-project.org/) following established methods.

### 2.8 Differential metabolite analysis

OPLS-DA, a supervised multivariate method, was utilized to identify differentially accumulated metabolites (DAMs) between resistant and susceptible varieties. Metabolites with VIP ≥ 1 and FC ≥ 2 or ≤ 0.5 were considered significantly differential. Selected compounds were characterized using the KEGG database (http://www.kegg.jp/kegg/pathway.html)^[25]^. Pathway enrichment was evaluated against the background set using a hypergeometric test, with significance defined ats P < 0.05.

## Discussion

Chestnut is a nutritious food crop, but its production is threatened by several pests, particularly the red spider mite (*O. ununguis*), which severely damages leaves by sucking plant sap. Developing mite-resistant chestnut varieties would reduce insecticide dependence and support a more sustainable, “green” chestnut industry. To date, most studies have focused on chestnut quality and yield, while little progress has been made in identifying germplasm resources with resistance-related molecules. This lack of knowledge limits the development of mite resistant crops through the use of molecular breeding. In this study, four chestnut germplasms with different resistance levels were analyzed under red spider mite stress. Metabolomic approaches were applied to identify key pathways and resistance-associated metabolites.

Metabolomics is a powerful approach for studying plant growth and adaptation in challenging environments and has been widely applied to investigate responses to biotic stress. However, metabolic defenses of Chinese chestnut (*C. mollissima*) during *O. ununguis* infestation remain poorly characterized. In this study, 71 DAMs were mainly enriched in flavone/flavonol biosynthesis and phenylpropanoid biosynthesis pathways. The phenylpropanoid pathway contributes to cell wall reinforcement through lignin biosynthesis^[26-27]^, and enhanced secondary wall formation is known to improve resistance to both biotic and abiotic stress^[28-34]^. Flavone and flavonol biosynthesis also represent classic defense pathways, producing flavonoids such as quercetin and anthocyanins with antioxidant, antimicrobial, and natural enemy-attracting functions. After pest (mite) feeding, trees can rapidly accumulate stress-induced secondary metabolites, increasing defensive compounds that either poison the pests or attract their natural enemies^[35]^. For example, *Castanea henryi* shows an induced resistance response to chestnut gall wasp feeding, with changes in tannin content playing a significant role in defense^[36]^.

In this study, 27 DAMs were identified through Venn diagram analysis. Compounds with high accumulation in the resistant group included cinnamic acid, methyl caffeate, baicalein, gallocatechin, dehydrodiisoeugenol, and lucolipin, suggesting their involvement in chestnut resistance to red spider mite. Cinnamic acid, a key intermediate in the phenylpropanoid pathway, contributes to cell wall lignification and strengthens tissue barriers; it has also been linked to soybean resistance against *Cotinomyces nuclealis* and shown repellant effects on poplar longhorn beetles^[37-38]^. Baicalein exhibits anti-inflammatory, antioxidant, and antiviral activities and can inhibit pest digestive enzymes, making it a potential natural insecticide^[39-40]^. Gallocatechin, a precursor of condensed tannins, is important in chemical defense by disrupting insect feeding, digestion, and metabolism^[41]^. Dehydrodiisoeugenol, an intermediate of lignin biosynthesis, enhances cell wall fortification and may also interfere with insect digestive enzymes or induce oxidative stress in pathogens^[42]^. Luciferin is a triterpenoid that can directly inhibit insect feeding^[43]^, scavenge reactive oxygen species (ROS), and activate defense-related genes. The synthesis and regulation of such compounds reflect the multi-level defense strategies plants have developed through long-term evolution and provide a theoretical basis for green pesticide development and breeding of stress-resistant fruit trees.

The metabolic response of plants to spider mites is a complex process involving both the synthesis of specific metabolites^[44-45]^ and the regulation of hormone signaling networks^[46]^. At the primary defense level, chestnut leaves appear to form a chemical barrier against mite feeding by activating phenylpropanoid and flavonoid biosynthetic pathways. Several resistance markers (e.g., cinnamic acid, methyl caffeate, baicalin, gallocatechin, dehydrodiisoeugenol, and lucolipin) were found to accumulate specifically in highly resistant germplasm under mite stress, suggesting their role in resistance. However, metabolite changes alone cannot fully explain the mechanisms of resistance. Further studies on metabolite function and associated protein changes are needed to clarify the molecular basis of chestnut responses to mite stress. Overall, this study addressed the gap in knowledge of chestnut-mite interactions and provides valuable insights for elucidating anti-mite mechanisms, developing green biopesticides, and advancing insect-resistant germplasm breeding. Specifically, our results showed that chestnut germplasms with different resistance levels exhibit distinct metabolic responses

## Conclusion

This study investigated metabolic changes in Chinese chestnut leaves after red spider mite feeding, comparing resistant and susceptible varieties. It represents the first metabolomics report in Chinese chestnut, identifying 713 metabolites in total. In comparisons of the resistant variety (R-1) with susceptible varieties (M1, M2, and S19), 128, 133, and 118 DAMs were detected, respectively. These DAMs were found to be related to pathways such as phenylalanine metabolism, flavonoid biosynthesis, anthocyanin biosynthesis, and D-amino acid metabolism. The findings indicate that phenylpropanoids and flavonoids play central roles in resistance, providing new insight into chestnut-mite interactions and a basis for breeding mite-resistant chestnut varieties.

## Acknowledgements

The authors would like to thank TopEdit (www.topeditsci.com) for its linguistic assistance during the preparation of this manuscript.

## Author Contributions

XFZ, SHZ, YL, YG, JYL, LYF, YJW, and GPW contributed to the study conception and design. Material preparation, data collection, and analysis were performed by XFZ and YJW. The first draft of the manuscript was written by XFZ. GPW commented on previous versions of the manuscript. All authors read and approved the final manuscript.

## Availability of data and materials

Access to the electronic supplementary material were presented in Table S1, Figure S2 and Figure S3.

## Ethical approval

This research did not utilize either animals or human participants.

